# BNIP3-mTOR Signaling Mediates Resistance to MET Inhibition in Glioblastoma

**DOI:** 10.64898/2025.12.28.696781

**Authors:** Yunzhan Li, Hanif Khan, Seyma Demirsoy, William Bernhardt, Hannah Valensi, Jeongwu Lee, Mitchell Machtay, Dawit Aregawi, Michael Glantz, Pierre Giglio, Shengyu Yang, Todd Schell, Vonn Walter, Yasin Uzun, Inan Olmez

## Abstract

Glioblastoma (GBM) is an aggressive primary brain malignancy with poor prognosis due to rapid progression, extensive invasiveness, and intrinsic resistance to standard therapies. Aberrant activation of receptor tyrosine kinases (RTKs), particularly MET, drives tumor proliferation, invasion, and therapy resistance. Here, we show that MET inhibition with crizotinib induces senescence and mitochondrial dysfunction in glioma-initiating cells (GICs), in part via downregulation of the mitochondrial protein BNIP3. However, BNIP3 downregulation activates mTOR signaling, enabling adaptive resistance. Targeting mTOR with everolimus in combination with crizotinib synergistically enhances anti-tumor effects, inducing apoptosis, senescence, and necroptosis, and significantly reducing cell viability and sphere-forming capacity. In orthotopic GBM xenograft models, this combination, particularly in a sequential regimen, markedly prolongs survival without overt toxicity. Our findings identify a BNIP3–mTOR signaling axis as a critical mediator of resistance to MET inhibition and provide a mechanistic rationale for combined MET and mTOR targeting as a promising therapeutic strategy in GBM.

**Statement of Translational Relevance:** Glioblastoma (GBM) remains a highly aggressive and treatment-resistant brain tumor with limited therapeutic options. Our study identifies a BNIP3-mTOR signaling axis as a key mediator of resistance to MET inhibition. We show for the first time that combined MET and mTOR inhibition exhibits synergistic effects against GBM in vitro and in vivo. This combination prolongs survival without overt toxicity, providing a strong preclinical rationale for clinical evaluation in GBM patients with high MET expression and offering a promising strategy to overcome adaptive resistance.

## INTRODUCTION

Glioblastoma (GBM) is the most common and most lethal primary brain malignancy (1). It is characterized by extensive infiltration into surrounding brain parenchyma, rapid proliferation, and profound resistance to conventional therapies. Despite multimodal treatment consisting of maximal surgical resection followed by radiotherapy and temozolomide chemotherapy, patient outcomes remain dismal, with a median survival of only 15–18 months following diagnosis (2), underscoring the urgent need for new therapies. Receptor tyrosine kinase (RTK) signaling pathways are dysregulated in 80-90% of GBMs through genetic and non-genetic modifications (3,4). These signaling abnormalities play a central role in tumor growth, invasion, and therapeutic resistance, and have therefore positioned RTKs as attractive targets for GBM therapy. However, despite extensive clinical evaluation of RTK inhibitors, durable clinical benefit has remained elusive.

Among RTKs, the hepatocyte growth factor (HGF)/MET signaling axis has emerged as a particularly compelling therapeutic target in GBM. Activation of MET triggers multiple downstream pathways, including PI3K/AKT, RAS/MAPK, and STAT3, which collectively promote tumor cell survival, proliferation, invasion, and the maintenance of glioma stem-like populations (5–8). Aberrant MET activation is frequently observed in GBM and is particularly enriched at invasive tumor margins, where it contributes to the highly infiltrative and aggressive phenotype of this tumor (9). Moreover, MET signaling has been directly implicated in therapeutic resistance by enhancing DNA damage repair following irradiation and promoting tumor cell survival under hypoxic stress, thereby diminishing the efficacy of standard treatments such as radiotherapy and temozolomide (10). These properties have driven substantial interest in MET as a therapeutic target in GBM.

Accordingly, several MET inhibitors, including crizotinib and cabozantinib, have been evaluated in preclinical GBM models and early-phase clinical trials, with mixed therapeutic outcomes (9,11–13). While MET-directed therapies can elicit initial anti-tumor responses, their clinical efficacy as monotherapies has been limited and transient, largely due to the rapid development of resistance (14,15). Resistance to MET-targeted monotherapy arises through adaptive signaling rewiring and engagement of compensatory survival pathways that sustain downstream oncogenic signaling despite effective MET inhibition (16–18). However, the molecular mechanisms governing this adaptive resistance remain incompletely understood.

In this study, we identify a previously unrecognized mechanism underlying resistance to MET inhibition in GBM. We demonstrate that MET inhibitor crizotinib exhibits anti-tumor activity, at least in part, through suppression of BNIP3, a mitochondrial protein that regulates mitophagy and cell death. We further show that BNIP3 downregulation results in activation of mTOR signaling, thereby promoting adaptive survival responses following MET inhibition. Importantly, pharmacologic inhibition of mTOR with everolimus markedly enhances the anti-tumor efficacy of crizotinib both in vitro and in vivo. Together, our findings uncover a BNIP3–mTOR signaling axis as a critical determinant of MET inhibitor response and provide a strong rationale for combinatorial targeting of MET and mTOR as a therapeutic strategy in GBM.

## MATERIALS AND METHODS

### Cell culture, cell viability detection, self-renewal assays, and reagents

Primary glioma-initiating cell (GIC) lines were obtained from University of Alabama and Jeongwu Lee (Cleveland Clinic) and have been published previously (19–21). The human identity of all cell lines was verified by short tandem repeat (STR) profiling prior to experimentation. GIC identity was further confirmed by the expression of established stem cell markers, including SOX2, CD133, OLIG2, and CD44. All cell lines were routinely tested for mycoplasma contamination by PCR, with testing repeated every five weeks. To minimize genetic drift and ensure experimental reproducibility, all experiments were performed using low-passage cells (<10 passages). Cultures were periodically reinitiated from early-passage stocks every four weeks. GICs were maintained as floating neurospheres in Neurobasal medium supplemented with glutamine, N2 (ThermoFisher Scientific), B27 (ThermoFisher Scientific), epidermal growth factor (EGF; 20 ng/mL; R&D Systems), and fibroblast growth factor (FGF; 20 ng/mL; R&D Systems) at 37°C in a humidified incubator with 5% CO₂. For in vitro assays, neurospheres were dissociated into single cells using 0.02% ethylenediaminetetraacetic acid (EDTA; Lonza) and plated onto laminin-coated culture plates (Corning). Self-renewal assays were conducted using ultra-low attachment plates (Corning). Neurosphere formation was quantified following drug treatment, and analyses were performed as previously described (22). Cell viability was assessed 72 hours after treatment using both the CellTiter-Glo luminescent assay (Promega) and Trypan Blue exclusion with automated cell counting (Cellometer Auto T4; Nexcelom). Crizotinib (HY-50878) was purchased from MedChemExpress, and everolimus (S1120) was obtained from Selleckchem.

### Animal studies

All animal studies were approved by the Institutional Animal Care and Use Committee (IACUC) at Penn State University and conducted in accordance with institutional guidelines. Six-to-eight-week-old female BALB/c SCID NCr mice were used for all in vivo experiments. A total of 150,000 G827 or G559 GICs were stereotactically injected into the right striatum of each mouse. Following tumor implantation, mice were randomized into four treatment groups: vehicle control, crizotinib alone, everolimus alone, or combination treatment. Two treatment regimens were employed: (i) a continuous treatment regimen, in which crizotinib and everolimus were administered daily throughout the study; and (ii) a sequential combination regimen (Combo 2), in which crizotinib alone was administered daily for the first 10 days and continued thereafter, with daily everolimus added beginning on day 11. Both drugs were administered once daily by oral gavage. Crizotinib and everolimus were dosed at 50 mg/kg and 5 mg/kg, respectively. Mice were monitored daily for neurological symptoms and overall health and were euthanized upon reaching predefined humane endpoints or terminal disease. Overall survival was recorded and compared between treatment groups. No animals were excluded from the survival analysis. Sample size for each group was determined based on prior experience and power calculations, assuming a median survival of 40 days for the control group and 70 days for treated groups, with a maximum follow-up period of 120 days and a statistical power of 85%.

### Immunoblotting, Caspase-3/7 assay, and qPCR

Immunoblotting was performed as previously described (23). The following primary antibodies were used for immunoblotting: β-actin (A5441) and GAPDH (G9545) were obtained from Sigma-Aldrich; phospho-mTOR (Ser2448; #2971), phospho-p70 S6 kinase (Thr389; #9234), phospho-S6 (Ser235/236; #4858), phospho-RIP1 (#65746), BNIP3 (#44060), and H3K27me3 (#9733) were obtained from Cell Signaling Technology; H3K9me2 (ab1220) was obtained from Abcam; histone H3 (05–499) was obtained from MilliporeSigma; and phospho-MLKL (PA5-105678) and cleaved caspase-9 (PA5-105271) were obtained from Thermo Fisher Scientific. Caspase-3/7 activity was measured using the Caspase-Glo® 3/7 Assay kit (G8090; Promega) according to the manufacturer’s instructions following 48 hours of drug treatment. Total RNA was isolated using QIAzol reagent (Qiagen) and reverse-transcribed into cDNA using the SuperScript™ III First-Strand Synthesis Kit (Invitrogen), following the manufacturer’s instructions. Quantitative real-time PCR was performed using 2 µL of diluted cDNA on an Applied Biosystems StepOnePlus PCR system with Power SYBR™ Green PCR Master Mix (Applied Biosystems). Relative gene expression levels were calculated for each sample and normalized to *GAPDH* expression for comparative analysis.

The following primer sequences were used for qPCR (5′ → 3′):

- *IL6*: F: ACTCACCTCTTCAGAACGAATTG, R: CCATCTTTGGAAGGTTCAGGTTG
- *CXCL10*: F: GTGGCATTCAAGGAGTACCTC, R: TGATGGCCTTCGATTCTGGATT
- *CXCL8*: F: TTTTGCCAAGGAGTGCTAAAGA, R: AACCCTCTGCACCCAGTTTTC
- *BNIP3*: F: CAGGGCTCCTGGGTAGAACT, R: CTACTCCGTCCAGACTCATGC

### Plasmid and siRNA transfection

An mTOR expression plasmid (Addgene #69010), along with the corresponding control plasmid, was used for rescue experiments. A total of 125,000 cells were seeded onto laminin-coated 12-well plates, and plasmid transfection was performed the following day using FuGENE® HD transfection reagent (Promega) according to the manufacturer’s instructions. Twenty-four hours after transfection, cells were treated with a combination of crizotinib and everolimus. Cell numbers were quantified and compared after 48 hours of treatment. Non-targeting control siRNA and siRNAs targeting *c-MET* and *BNIP3* were obtained from Dharmacon (SMARTpool ON-TARGETplus). siRNA transfections in GICs were performed using Lipofectamine® RNAiMAX (Thermo Fisher Scientific) in accordance with the manufacturer’s protocol, with a final siRNA concentration of 10 nmol/L.

### Bulk RNA-seq and Bioinformatical analysis

Total RNA was extracted using the Qiagen RNeasy Kit (Qiagen), and sequencing libraries were constructed with the NEB Next Ultra RNA Library Prep Kit for Illumina platform sequencing. Paired-end RNA sequencing reads were aligned to the human reference genome (hg38) using the STAR aligner (24). Gene-level quantification was performed using Cufflinks (25) with the Gencode v38 annotation (26), and expression levels were reported as fragments per kilobase of transcript per million mapped reads (FPKM). Genes with an FPKM greater than 1.0 in at least two samples from either experimental group were retained for downstream analysis. Raw read counts were generated using the *featureCounts* function from the Rsubread package (27). Differential expression analysis was carried out using edgeR (28). Raw counts were normalized using the trimmed mean of M-values (TMM) method, and a quasi-likelihood negative binomial generalized log-linear model was fit using the glmQLFit and glmQLFTest functions. Genes with a false discovery rate (FDR) at or below 0.05 and an absolute log_2_ fold change at or above 1.5 were considered significantly differentially expressed. Gene set enrichment analysis (GSEA) was performed using the GSEA software suite (29) with normalized FPKM values as input. Visualization of differential expression and enrichment results was performed using R packages including *ggplot2*, *pheatmap*, *enrichplot*, and *dplyr*.

### Mitochondrial membrane potential analysis (JC-1 staining)

Mitochondrial membrane potential was assessed using JC-1 dye (Thermo Fisher Scientific) according to the manufacturer’s instructions. Briefly, GICs were seeded onto laminin-coated plates and treated as indicated. Following treatment, cells were incubated with JC-1 working solution at 37°C for 20 minutes in the dark. After incubation, cells were washed twice with assay buffer and analyzed immediately by fluorescence microscopy. JC-1 aggregates (red fluorescence) indicate healthy mitochondria with high membrane potential, whereas JC-1 monomers (green fluorescence) indicate depolarized mitochondria.

### Mitochondrial Stress Test

Mitochondrial Stress Test was performed as described previously (21). Cells were seeded as a monolayer onto a laminin-coated Seahorse 24-well tissue culture plate (Agilent Technologies, Santa Clara, CA). After 48 hours of drug treatment, the media was replaced with DMEM supplemented with pyruvate, and the cells were allowed to equilibrate for 30 minutes at 37°C in a CO₂-free incubator. Oxygen consumption rate (OCR) was measured using a Seahorse XF24 Flux Analyzer (Agilent Technologies, Santa Clara, CA). After three baseline measurements, compounds were sequentially injected: 1 µM Oligomycin (Sigma-Aldrich), 2 µM BAM15 (Cayman Chemical), and 1 µM Antimycin A & 100 nM Rotenone (Sigma-Aldrich). Basal respiration was calculated by subtracting the average OCR after Antimycin A & Rotenone treatment from the first three baseline measurements. Maximum respiratory capacity was determined by subtracting the post-BAM15 OCR from the post-Antimycin A & Rotenone OCR. Reserve capacity was calculated by subtracting the basal OCR from the post-BAM15 OCR.

### β-Galactosidase and immunohistochemistry (IHC) staining

Senescence-associated β-galactosidase (SA-β-Gal) activity was assessed using a β-Gal staining kit (Abcam) according to the manufacturer’s instructions. Briefly, GICs were seeded onto laminin-coated plates and treated as indicated. After treatment, cells were washed with PBS and fixed using the provided fixation solution for 10–15 minutes at room temperature. Following fixation, cells were incubated with β-Gal staining solution at 37°C (without CO₂) overnight. Blue-stained cells were visualized and quantified using bright-field microscopy. IHC staining was performed as described previously (30). Tissue slides were fixed in a 1:1 mixture of acetone and methanol for 10 minutes at room temperature. Endogenous peroxidase activity was quenched using peroxidase inhibitor (CM1; Ventana) for 8 minutes. Slides were then incubated with primary antibodies at room temperature. Antigen–antibody complexes were subsequently detected using the DISCOVERY OmniMap Anti-Rabbit HRP detection system in combination with the DISCOVERY ChromoMap DAB Kit (Ventana Medical Systems), following the manufacturer’s instructions. Stained slides were visualized by bright-field microscopy.

### Statistics and synergy calculations

All statistical analyses were performed using GraphPad Prism 10 (GraphPad Software). For comparisons between two groups, Student’s t-test was applied, while one-way ANOVA followed by Tukey’s post-hoc test was used for multiple group comparisons. P-values <0.05 were considered statistically significant (α = 0.05). Kaplan–Meier survival analysis was employed to generate mouse survival curves. Synergy between treatments was evaluated using two independent methods: the Bliss independence model and the Chou–Talalay method (ComboSyn). Combination indices (CI) were calculated according to the Chou–Talalay method, with CI < 1 indicating synergy and CI < 0.2 indicating strong synergy (31). The Bliss difference was calculated as described previously (32). Briefly, the Bliss value was obtained by subtracting the predicted cytotoxicity, based on individual treatments, from the observed cytotoxicity of the combination therapy. A Bliss value of zero corresponds to an additive effect, a value greater than zero indicates synergy, and a value less than zero indicates antagonism. This approach provides informative synergy assessment even when one component of a combination exhibits minimal effect.

## RESULTS

### MET expression is elevated in GBM and correlates with poor patient survival

We first assessed MET expression in patient GBM samples by IHC and observed high MET protein levels, as quantified by H-score, in GBM specimens (Fig. 1A). To evaluate the clinical relevance of MET expression, we analyzed The Cancer Genome Atlas (TCGA) GBM datasets and found that increased MET expression was associated with significantly shorter overall survival (Fig. 1B). Consistent with these findings, analysis across glioma grades within the TCGA cohort revealed a progressive increase in MET expression with higher tumor grade, indicating a positive correlation between MET expression and glioma malignancy (Fig. 1C). GBM has been classified into four molecular subtypes—classical, proneural, neural, and mesenchymal—based on gene expression profiling (33,34). In epithelial malignancies, cancer cells acquire mesenchymal characteristics through epithelial–mesenchymal transition (EMT), a process associated with therapeutic resistance, enhanced invasiveness, and metastatic potential (35). Similarly, acquisition of mesenchymal features in GBM is linked to increased tumor aggressiveness and resistance to therapy, underscoring the need for effective strategies targeting the mesenchymal phenotype. Analysis of TCGA datasets showed that MET mRNA expression was significantly elevated in the mesenchymal subtype compared with other GBM subtypes (Fig. 1D). Consistently, MET expression was enriched in infiltrating tumor regions, the leading tumor edge, and pseudopalisading cells, indicating an association with aggressive tumor features (Fig. 1E).

**Figure 1.**
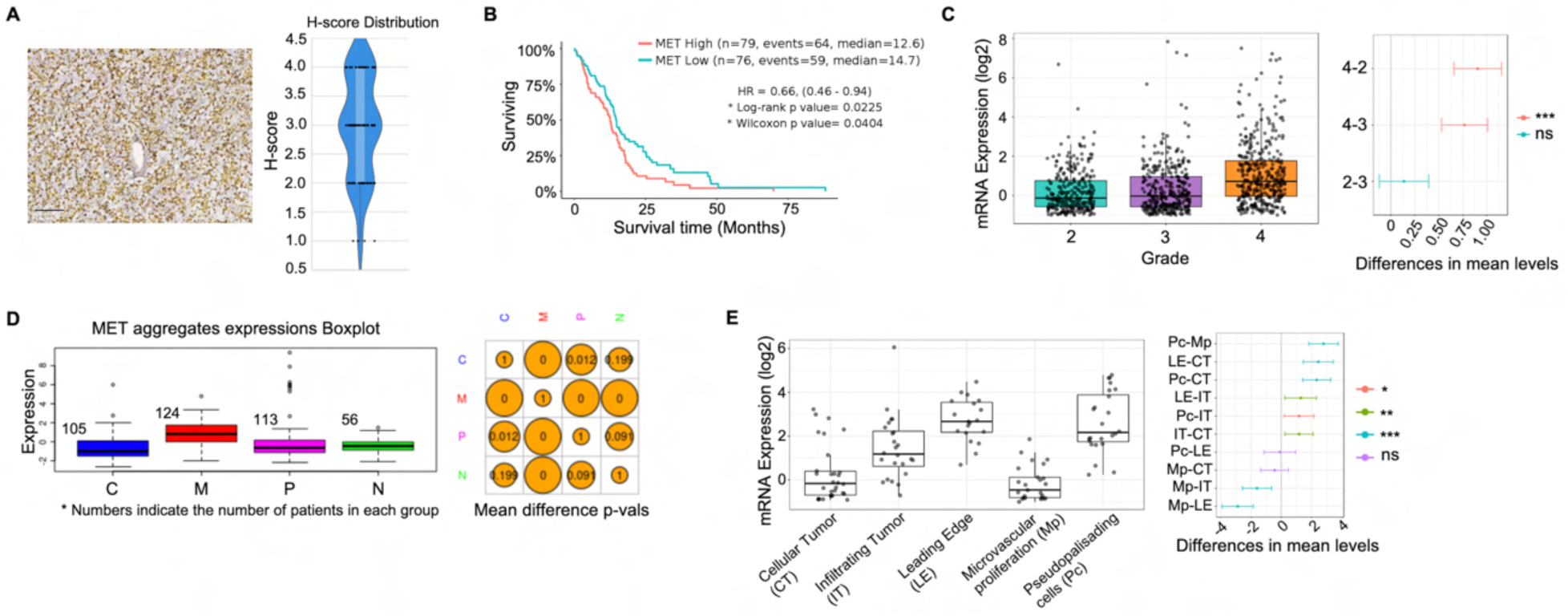
MET expression is elevated in glioblastoma and associates with tumor aggressiveness and poor survival. (A) Representative immunohistochemical staining of MET in human GBM tissue (left) and quantification of MET protein expression using H-score analysis (right), demonstrating high MET expression across GBM samples. n = 28 GBM samples (from Ivy GBM Atlas Project). Scale bar, 100 μm. (B) Kaplan–Meier survival analysis of GBM patients from the TCGA database stratified by MET expression levels, showing significantly reduced survival in patients with high MET expression. Hazard ratio (HR), log-rank, and Wilcoxon p values are indicated. (C) MET mRNA expression across glioma grades (II–IV) in the TCGA dataset, revealing a progressive increase in MET expression with advancing tumor grade. Right panel shows pairwise comparisons of mean expression differences between grades. (D) MET mRNA expression across molecular GBM subtypes—classical (C), mesenchymal (M), proneural (P), and neural (N)—demonstrating significant enrichment of MET expression in the mesenchymal subtype. Numbers indicate patient counts per group; right panel shows pairwise mean difference p values. (E) MET mRNA expression across distinct anatomic tumor regions, including cellular tumor (CT), infiltrating tumor (IT), leading edge (LE), microvascular proliferation (Mp), and pseudopalisading cells (Pc), showing preferential enrichment of MET expression in invasive and aggressive tumor compartments. Right panel depicts pairwise comparisons of mean expression differences.

### Prolonged MET inhibition induces senescence and alters transcriptional programs in GBM

To investigate the transcriptional response to MET inhibition, glioma-initiating cells (GICs) were treated with the MET inhibitor crizotinib for three cycles (3 days per cycle). On day 10, surviving clones were collected and subjected to bulk RNA sequencing alongside untreated control cells (Fig. 2A). RNA-seq analysis revealed significant upregulation of genes associated with p53 signaling and cellular senescence in crizotinib-treated GICs (Fig. 2B). GSEA and pathway analysis showed downregulation of DNA synthesis, cell cycle progression, and mitosis compared with controls (Fig. 2C and Supplementary Fig. 1A). Senescence induction was validated by β-Galactosidase staining, which showed a marked increase in senescent cells after prolonged crizotinib treatment (Fig. 2D, Supplementary Fig. 1B). Consistently, p53 protein levels were elevated (Fig. 2E), and mRNA expression of senescence-associated cytokines IL-6, CXCL10, and CXCL8 was increased (Fig. 2F, Supplementary Fig. 1C). GSEA further revealed enrichment of DNA methylation and histone lysine methylation pathways. Immunoblot analysis confirmed increased histone H3 modifications, including H3K9me2/3 and H3K27me3, as well as total H3 levels, in crizotinib-treated GICs (Supplementary Fig. 1D–E). Collectively, these results demonstrate that prolonged MET inhibition triggers a senescence program in GICs while suppressing proliferation- and metabolism-related pathways. Finally, we investigated the stability of the senescent state following MET inhibition. After withdrawal of crizotinib, GICs initially exhibited slow proliferation consistent with senescence, but resumed growth over time, albeit at a slower rate than untreated controls. Similarly, in an orthotopic brain tumor model, crizotinib-treated GICs prolonged survival compared with controls, but the difference was modest, indicating that the senescent state is not permanent (Fig. 2G).

**Figure 2.**
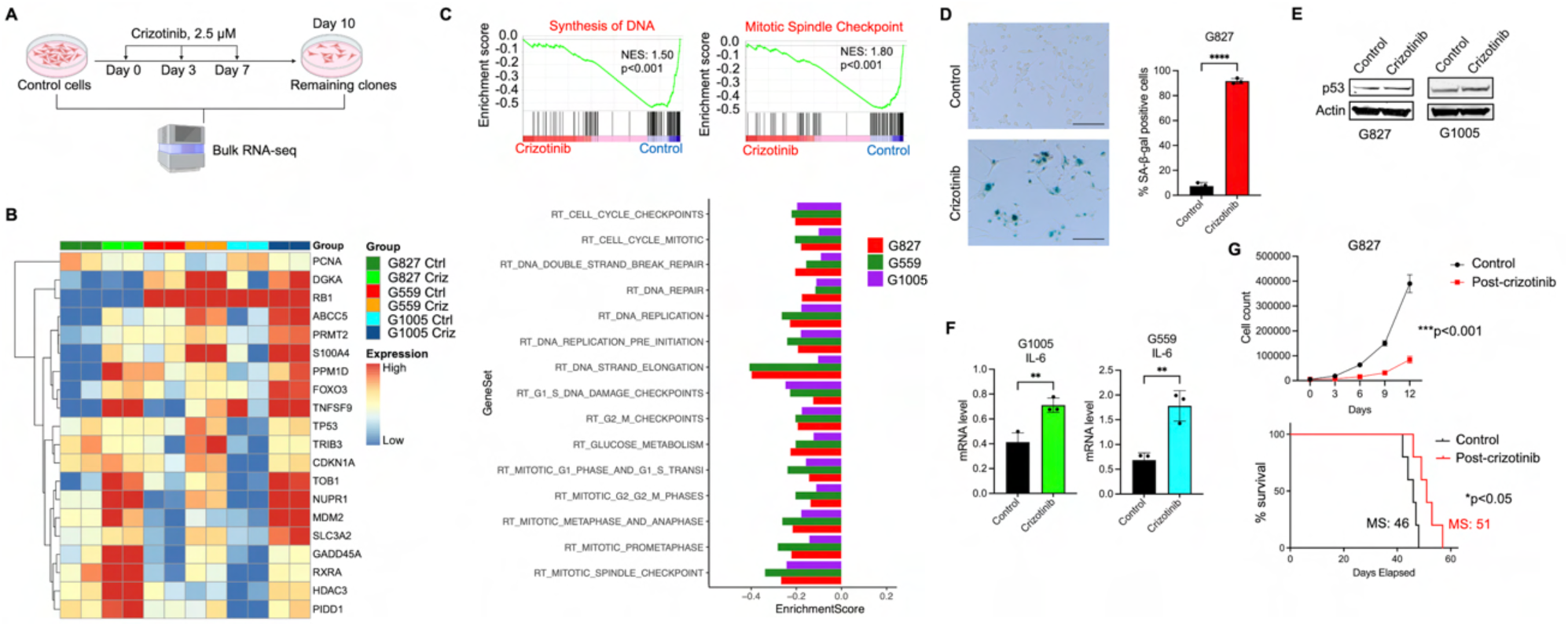
MET inhibition induces a senescence-like state and suppresses proliferative programs in GICs. (A) Experimental design: GICs were treated with crizotinib for three cycles (3 days per cycle), and surviving clones were collected for bulk RNA sequencing alongside controls. (B) Heatmap of differentially expressed genes showing upregulation of p53 signaling and senescence-associated genes. (C) GSEA and Reactome pathway analysis demonstrating significant downregulation of DNA synthesis, mitosis, and cell cycle progression in crizotinib-treated cells. Normalized enrichment scores (NES) and p values are indicated. (D) Representative images of senescence-associated β-galactosidase (SA-β-gal) staining in control and crizotinib-treated GICs, showing increased senescent cells upon MET inhibition. Crizotinib (1.5 µM). Scale bar, 200 µm. Data are shown as mean ± SEM. (****p < 0.0001; two-tailed *t*-test). (E) Immunoblot analysis showing increased p53 protein levels in crizotinib-treated cells. (F) Quantitative RT–PCR analysis of IL-6 mRNA expression in GICs following crizotinib (1.5 µM) treatment, indicating induction of a senescence-associated secretory phenotype (SASP). Data are shown as mean ± SEM. (**p < 0.01; two-tailed *t*-test). **(G)** Upper panel: Cell proliferation curves showing reduced growth of GICs following prolonged crizotinib treatment (two-tailed *t*-test). Lower panel: Kaplan–Meier survival analysis demonstrating extended survival in mice implanted with crizotinib-pretreated GICs compared with controls. Median survival (MS) values and p values are indicated.

To further dissect and validate the tumor-suppressive transcriptional program observed in Figure 2, we performed pathway-level analysis using Ingenuity Pathway Analysis (IPA). IPA confirmed key pathway alterations, including activation of senescence- and p53-associated programs and suppression of cell cycle–related processes, and additionally revealed downregulation of HIF1α-regulated hypoxia signaling in crizotinib-treated GICs (Fig. 3A). Among the most significantly downregulated transcripts, we identified BNIP3, a canonical HIF1α target gene involved in hypoxic stress adaptation (Fig. 3B). Analysis of the TCGA GBM dataset further showed that BNIP3 expression positively correlates with other HIF1α-regulated genes (Supplementary Fig. 2A). Consistent with the RNA-seq data, BNIP3 protein levels were markedly decreased in crizotinib-treated tumors compared with controls, as confirmed by immunoblotting and IHC (Fig. 3B). Using H-score quantification, we observed high BNIP3 expression in GBM patient samples (Supplementary Fig. 2B), consistent with its role as a HIF1α-regulated gene and highlighting its potential contribution to GBM pathophysiology. To assess the functional role of BNIP3, we silenced it using siRNA, which significantly reduced cell viability and sphere-forming capacity in GICs (Fig. 3C and Supplementary Fig. 2C). BNIP3 downregulation in crizotinib-treated GICs prompted us to examine its impact on mitochondrial function, as BNIP3 localizes to mitochondria and promotes mitophagy (Fig. 3D). Pathway analysis using IPA revealed that prolonged crizotinib treatment was associated with increased mitochondrial dysfunction (Supplementary Fig. 2D), suggesting that BNIP3 suppression contributes to impaired mitochondrial homeostasis. Consistently, Seahorse analysis demonstrated that crizotinib-treated GICs exhibited increased oxygen consumption rate (OCR), indicative of altered mitochondrial activity (Fig. 3D). Since the increase in mitochondrial activity observed upon crizotinib treatment could not be fully explained by BNIP3 downregulation alone, we investigated potential adaptive resistance mechanisms. BNIP3 has been reported to interact with Rheb and suppress mTOR signaling (36) (Supplementary Fig. 2E), leading us to hypothesize that crizotinib-induced BNIP3 downregulation might result in enhanced mTOR activity. To test this, BNIP3 was silenced using siRNA, which led to activation of the mTOR pathway as demonstrated by immunoblot analysis (Fig. 3E). In contrast, pharmacologic inhibition of mTOR with everolimus reduced BNIP3 expression, whereas mTOR overexpression using an mTOR plasmid significantly elevated BNIP3 levels (Fig. 3F). Together, these findings support a bidirectional regulatory relationship between BNIP3 and mTOR signaling in GICs. Moreover, time-course immunoblot analysis following crizotinib treatment revealed an inverse correlation between BNIP3 expression and mTOR activity, with prolonged treatment resulting in progressive BNIP3 downregulation concomitant with increased mTOR activation (Fig. 3G). MET silencing using siRNA similarly induced mTOR activation (Supplementary Fig. 2F), further highlighting the interplay between BNIP3, mTOR signaling, and MET in adaptive resistance mechanisms.

**Figure 3.**
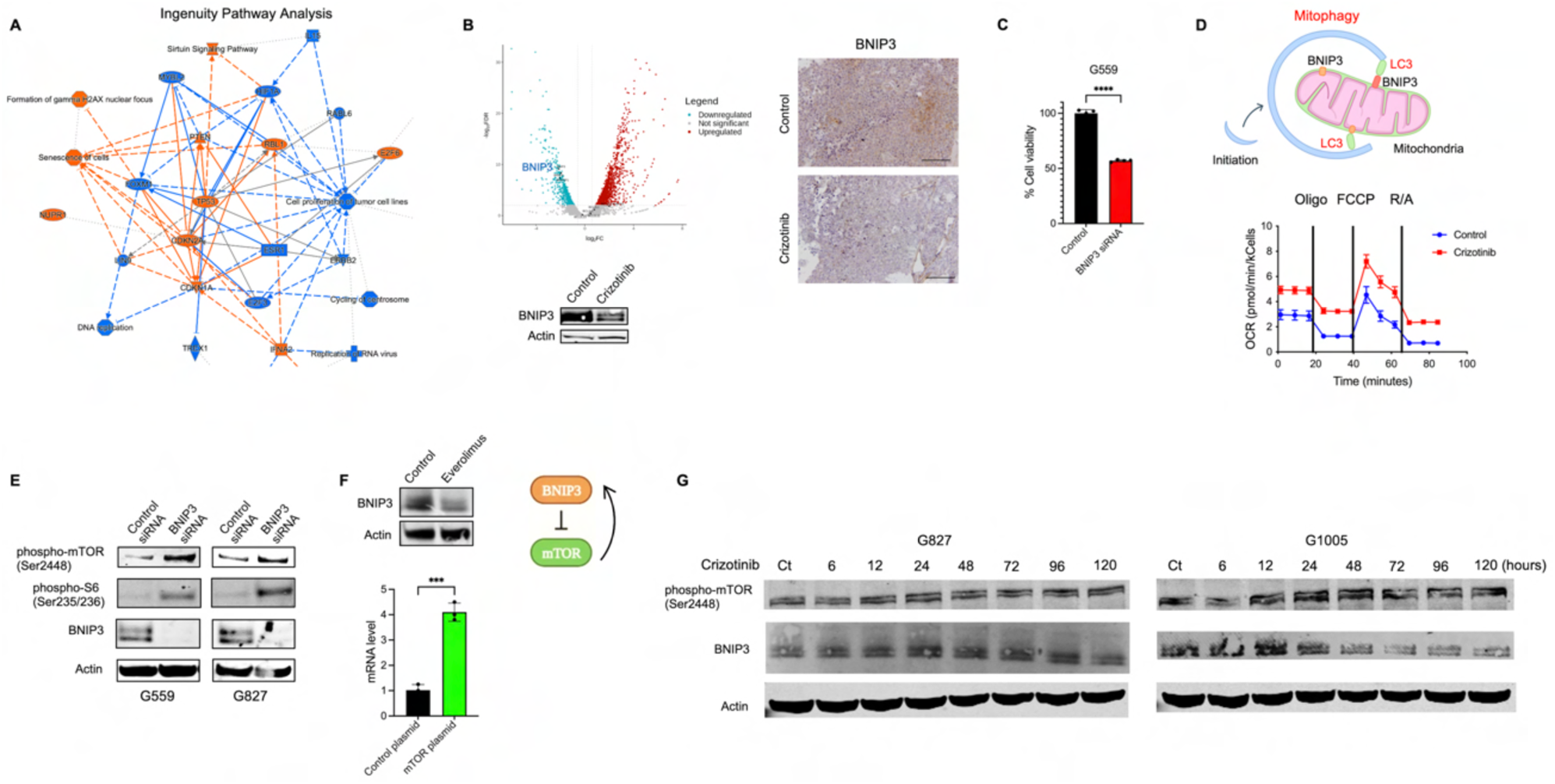
BNIP3 downregulation links MET inhibition to mitochondrial dysfunction and mTOR activation. (A) Ingenuity Pathway Analysis (IPA) network generated from bulk RNA-seq data of crizotinib-treated GICs. (B) Left: Volcano plot showing differentially expressed genes following crizotinib treatment, with BNIP3 significantly downregulated. Right: Representative immunohistochemical staining of BNIP3 in orthotopic GBM mouse xenografts, comparing control and crizotinib-treated groups, demonstrating reduced BNIP3 expression upon MET inhibition. Scale bar, 50 µm. Lower panel: Immunoblot analysis confirming BNIP3 downregulation following crizotinib treatment in GICs. (C) Cell viability assay following BNIP3 silencing by siRNA, demonstrating a significant reduction in cell viability compared with control siRNA. Data are shown as mean ± SEM (****p < 0.0001; two-tailed *t*-test). (D) Upper panel: Schematic illustrating the role of BNIP3 in mitophagy through LC3-mediated mitochondrial sequestration. Lower panel: Oxygen consumption rate (OCR) and the maximal OCR were measured using a Seahorse XF 24 Flux Analyzer in GICs made adherent as per Methods and subjected to a mitochondrial stress test. (\*\**P* < 0.001; one-way ANOVA with post-hoc Tukey analysis. Values are mean ± SEM of triplicates). Crizotinib dose was 1.5 µM. (E) Immunoblot analysis demonstrating activation of mTOR signaling upon BNIP3 loss. (F) Upper panel: Immunoblot showing decreased BNIP3 protein levels following mTOR inhibition with everolimus. Lower panel: Quantitative RT–PCR analysis of BNIP3 mRNA expression following mTOR overexpression, indicating positive regulation of BNIP3 by mTOR signaling. Data are shown as mean ± SEM. (***p < 0.001; two-tailed *t*-test). Middle schematic summarizes the reciprocal regulatory relationship between BNIP3 and mTOR. (G) Time-course immunoblot analysis in GICs treated with crizotinib (1.5 µM) for the indicated time points, demonstrating progressive mTOR activation concomitant with BNIP3 downregulation during prolonged MET inhibition.

### Combining MET and mTOR inhibitors exhibits synergy against GICs

Given the activation of mTOR signaling following MET inhibition, we next evaluated whether pharmacologic mTOR inhibition could overcome this adaptive response and enhance the antitumor efficacy of crizotinib. Everolimus effectively suppressed crizotinib-induced mTOR activation and downstream signaling, as confirmed by immunoblotting (Fig. 4A and Supplementary Fig. 3A). Combined MET and mTOR inhibition resulted in a significantly greater reduction in GIC viability and sphere-forming capacity compared with either treatment alone, indicating that mTOR inhibition sensitizes GICs to crizotinib (Fig. 4B-C and Supplementary Fig. 3B-C). Consistently, caspase-3/7 activity assays demonstrated that the combination of crizotinib and everolimus significantly induced apoptosis, whereas apoptosis was minimal or absent following single-agent treatments (Fig. 4D).

**Figure 4:**
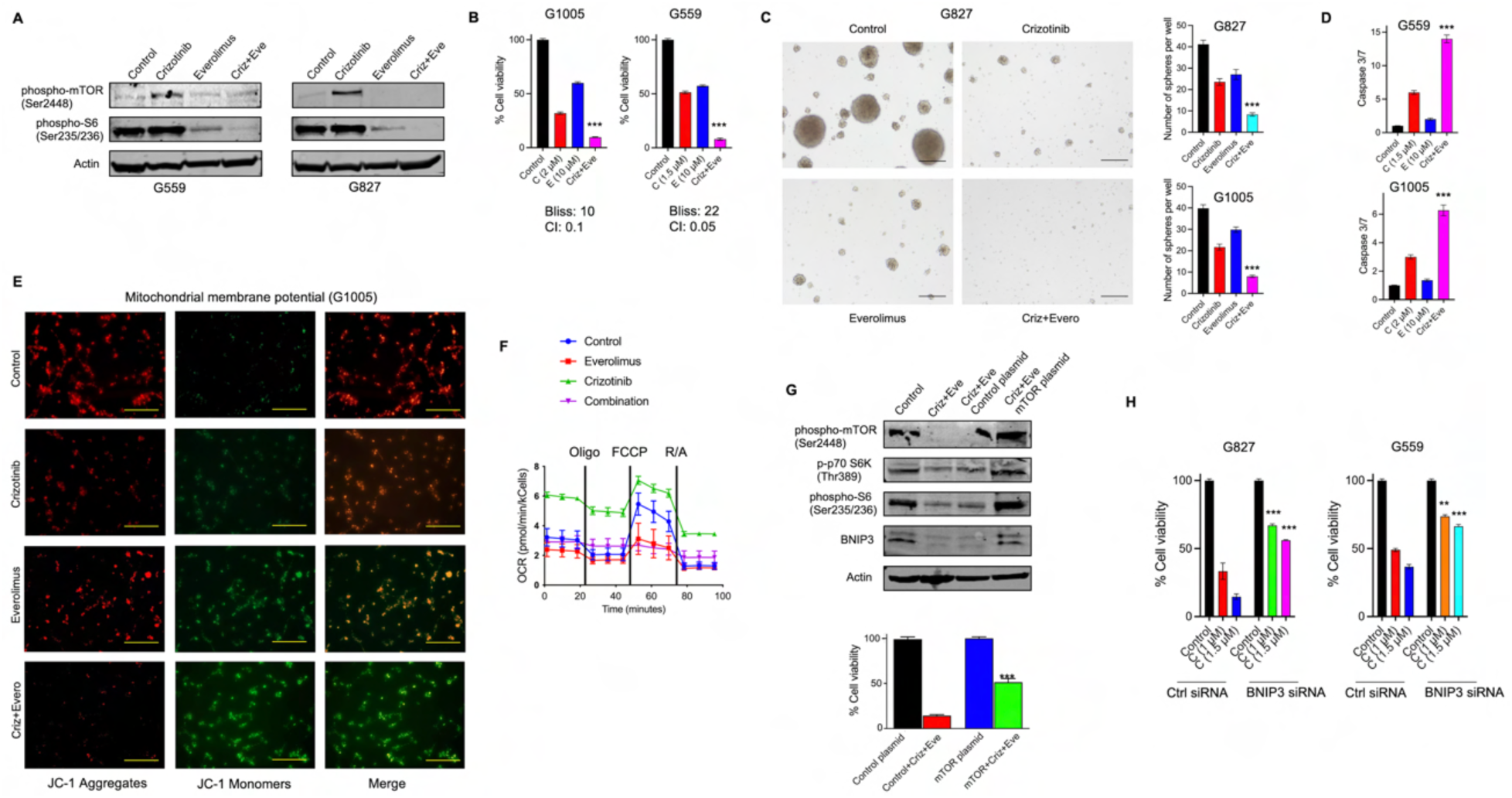
Combined MET and mTOR inhibition exhibits synergy against GICs. (A) Immunoblot analysis of mTOR signaling in GICs treated with crizotinib, everolimus, and their combination (Criz+Eve). (B) The combination of crizotinib (C) and everolimus (E) is synergistic against GICs (each treatment line received either DMSO or the indicated drug) (\*\*\**P* < 0.0001; one-way ANOVA with post-hoc Tukey analysis). Synergy was evaluated using Bliss independence and combination index (CI) analysis, indicating strong synergistic effects. (C) 100 GICs were cultured in 24-well plates over one week to compare sphere formation upon treatment with vehicle, crizotinib (1.5 µM), everolimus (5 µM), and the combination of crizotinib and everolimus (\*\*\**P* < 0.0001; one-way ANOVA with post-hoc Tukey analysis). Scale bars, 100 µm. (D) Caspase-3/7 level is increased with the three days of combined treatment. (\*\*\**P* < 0.0001; one-way ANOVA with post-hoc Tukey analysis). (E) Assessment of mitochondrial membrane potential in G1005 cells using JC-1 staining. Red fluorescence indicates JC-1 aggregates (intact membrane potential), whereas green fluorescence indicates JC-1 monomers (depolarized mitochondria). Scale bars, 200 µm. (F) Oxygen consumption rate (OCR) analysis measured by Seahorse assay in cells treated with everolimus, crizotinib, or the combination. The combination treatment reverses crizotinib-induced mitochondrial respiration. (G) Immunoblot analysis of mTOR signaling and BNIP3 in GICs transfected with control or mTOR plasmid and treated with Criz+Eve. Bottom, quantification of cell viability under the indicated conditions (\*\*\**P* < 0.0001; two-way ANOVA). (H) Cell viability of GICs transfected with control or BNIP3 siRNA and treated with crizotinib, indicating partial rescue of drug sensitivity upon BNIP3 depletion ((\*\**P* < 0.01, \*\*\**P* < 0.0001; two-way ANOVA).

Given the potential contribution of mTOR activation to the crizotinib-induced increase in mitochondrial activity, we next evaluated the effects of combined crizotinib and everolimus treatment on mitochondrial function. JC-1 staining revealed that combination treatment increased the proportion of monomeric JC-1, indicative of mitochondrial depolarization, while significantly reducing JC-1 aggregate formation (Fig. 4E and Supplementary Fig. 3D). Consistently, Seahorse OCR analysis demonstrated that combination treatment reversed the crizotinib-induced increase in mitochondrial respiration (Fig. 4F), collectively indicating that mTOR inhibition attenuates adaptive mitochondrial activation. Moreover, as shown in Supplementary Figure 3E, the combination therapy induced cellular senescence, evidenced by increased β-Galactosidase activity, further supporting the therapeutic efficacy of this approach. To directly assess the role of mTOR in mediating these effects, mTOR overexpression via plasmid largely restored BNIP3 levels and rescued cell viability in cells treated with the combination therapy, confirming that mTOR signaling regulates BNIP3 and modulates GIC sensitivity to crizotinib (Fig. 4G and Supplementary Fig. 3F). Finally, since crizotinib reduces BNIP3 expression in GICs, we investigated whether BNIP3 levels influence cellular response to crizotinib using BNIP3-specific siRNA. BNIP3 knockdown significantly decreased cellular sensitivity to crizotinib, suggesting that the anti-tumor effects of crizotinib are mediated, at least in part, through BNIP3 downregulation (Fig. 4H).

To further delineate the cell death mechanisms induced by the crizotinib and everolimus combination, we also evaluated apoptosis in GICs pre-treated with crizotinib. While combination treatment induced significant apoptosis in untreated parental cells (Fig. 4D), apoptosis levels were minimal in cells pre-treated with crizotinib for three cycles prior to everolimus administration, as detected by caspase-3/7 activity assay and immunoblot (Fig. 5A-B). Morphologically, these cells displayed necrotic features, suggesting a shift from apoptosis toward necroptosis (Fig. 5C). To investigate this further, we performed immunoblotting and tumor xenograft IHC analysis, which revealed increased expression of necroptosis markers following combined treatment (Fig. 5D-E). Collectively, these findings indicate that necroptosis substantially contributes to the cell death response elicited by the crizotinib and everolimus combination therapy.

**Figure 5.**
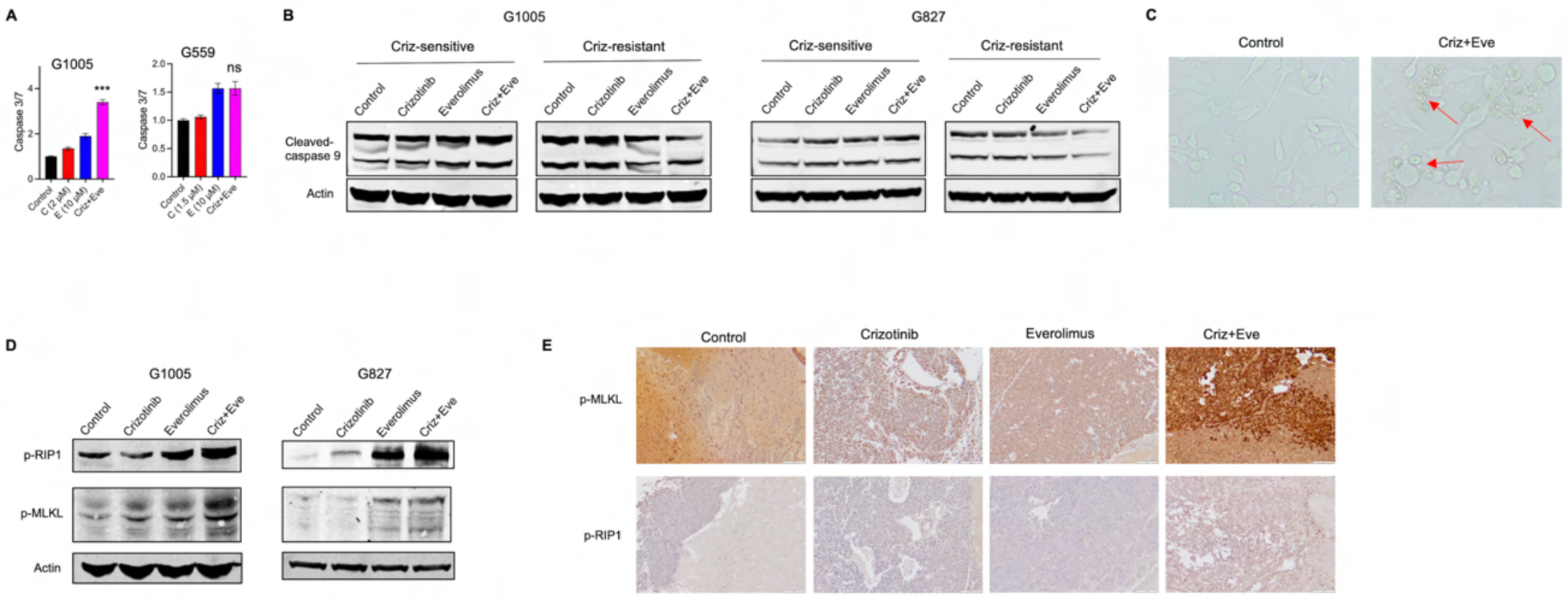
Combined crizotinib and everolimus treatment induces necroptotic cell death in crizotinib-pretreated GICs. (A) Caspase-3/7 activity in GICs treated with crizotinib (C), everolimus (E), or the combination (Criz+Eve). While a significant increase in apoptosis was observed in untreated cells, minimal apoptosis was detected in crizotinib-pre-treated cells (\*\*\**P* < 0.0001; one-way ANOVA, ns= non-significant). (B) Immunoblot analysis of cleaved caspase-9 in crizotinib-sensitive and crizotinib-pretreated GICs following treatment with crizotinib, everolimus, or their combination. (C) Representative phase-contrast images showing morphological changes consistent with cell death in cells treated with Criz+Eve compared with control. Arrows indicate rounded and detached cells. (D) Immunoblot analysis in crizotinib-pretreated GICs following the indicated treatments, indicating activation of necroptotic signaling upon combination therapy. (E) Immunohistochemical staining of orthotopic GBM mouse xenografts from control, crizotinib, everolimus, or Criz+Eve groups, demonstrating enhanced necroptosis in tumors receiving combination therapy. Scale bar, 50 µm.

### The combination of MET and mTOR inhibition is effective against an aggressive orthotopic GIC models in vivo

Next, we evaluated the efficacy of the combined crizotinib and everolimus treatment in orthotopic GBM models. For this purpose, we first used untreated parental GICs. Mice were treated with vehicle, once-daily oral crizotinib (50 mg/kg/day), once-daily oral everolimus (5 mg/kg/day), or the combination of both agents. While single-agent treatments produced minimal therapeutic effects, combination therapy significantly prolonged both median and overall survival (Fig. 6A). Given the distinct mechanisms of cell death observed between untreated and crizotinib-pretreated GICs, we next evaluated the efficacy of the combination therapy in crizotinib-pretreated GICs.

**Figure 6.**
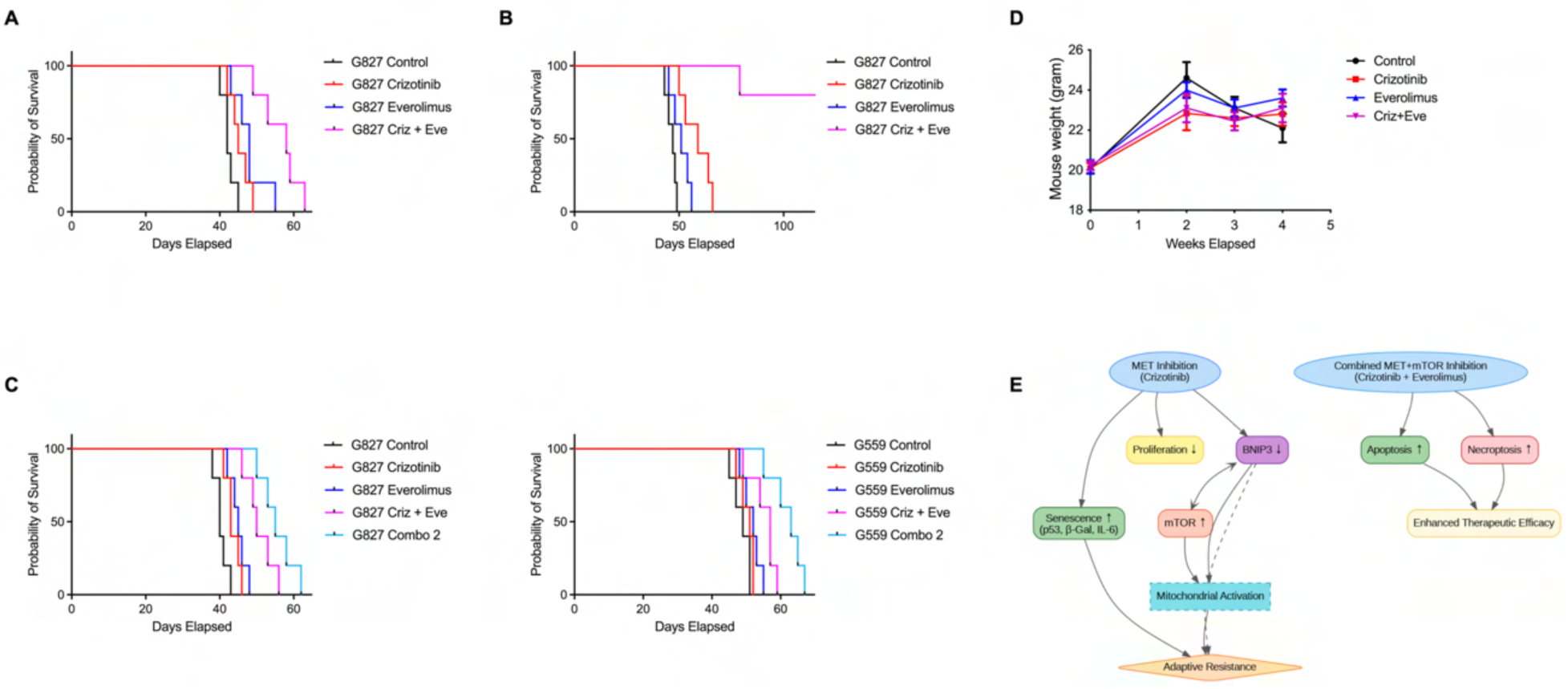
In vivo efficacy of combined MET and mTOR inhibition. Kaplan–Meier survival curves of mouse xenografts derived from (A) untreated parental GICs and (B) crizotinib-pretreated GICs. Mice received continuous daily treatment with crizotinib and everolimus throughout the study. (C) Kaplan–Meier survival curves of mouse xenografts from two GIC lines treated with (i) a continuous daily regimen of crizotinib and everolimus or (ii) a sequential combination regimen (Combo 2), in which crizotinib alone was administered daily for the first 10 days, followed by daily everolimus from day 11. (D) Mouse body weight comparison of each treatment group. (E) Schematic depicting the cooperation between everolimus and palbociclib.

After three cycles of in vitro crizotinib treatment, GICs were implanted intracranially to establish xenografts. Daily administration of crizotinib and everolimus markedly extended survival, with four of five mice surviving at day 150 (Fig. 6B). Building on these findings, we implemented a dosing regimen designed to mimic in vitro crizotinib treatment cycles, wherein crizotinib alone was administered daily for the first 10 days and continued thereafter, with daily everolimus added beginning on day 11 (Combo #2). This modified schedule further extended survival compared with the standard combination regimen (Fig. 6C). Finally, as a measure of toxicity, we followed mouse weights across all treatment groups. As shown in Figure 6D, mouse weights remained comparable across the control and treatment groups, with no overt toxicity observed following single-agent or combination therapy. Overall, our findings demonstrate that crizotinib and everolimus act synergistically through several mechanisms, resulting in significant antitumor efficacy against GICs (Fig. 6E).

## DISCUSSION

GBM remains one of the most aggressive and treatment-resistant human malignancies, characterized by rapid disease progression and dismal patient survival. Despite aggressive multimodal treatment approaches, tumor recurrence is inevitable, underscoring the urgent need for more effective and durable therapeutic strategies. In this context, our study identifies combined targeting of MET and mTOR signaling as a mechanistically informed approach to overcome adaptive resistance in GBM.

We demonstrate that MET expression is markedly elevated in GBM, particularly in the mesenchymal subtype, which is associated with increased invasiveness, therapeutic resistance, and poor patient outcomes (34,37). Consistent with these observations, pharmacologic inhibition of MET using crizotinib induces a robust senescent phenotype in GICs, accompanied by suppression of DNA synthesis, cell cycle progression, and proliferative capacity. However, the partial reversibility of this senescent state following crizotinib withdrawal indicates that MET inhibition alone is insufficient to induce durable tumor suppression, highlighting the need for rational combination strategies to prevent relapse and adaptive resistance.

A central finding of our study is the identification of a BNIP3–mTOR signaling axis as a critical mediator of resistance to MET inhibition. BNIP3, a HIF1α-regulated mitochondrial protein, plays a key role in mitophagy and cellular adaptation to metabolic and hypoxic stress (38,39). The downregulation of BNIP3 following MET inhibition suggests a potential mechanism underlying the observed mitochondrial dysfunction—a hallmark of metabolic reprogramming in cancer cells (40–42). We demonstrate that BNIP3 downregulation results in activation of mTOR signaling, thereby promoting tumor cell survival and adaptive resistance. The role of BNIP3 in regulating mitochondrial function through mTOR signaling is critically important, as mTOR is a master regulator of cellular metabolism and stress adaptation (43–45). This finding uncovers a key adaptive mechanism driving resistance to MET inhibition, offering a strong rationale for dual targeting of MET and mTOR.

We demonstrate that pharmacologic inhibition of mTOR with everolimus markedly enhances the anti-tumor effects of crizotinib. Combined treatment induces pronounced senescence, apoptosis, and mitochondrial dysfunction, resulting in reduced cell viability, impaired sphere-forming capacity, and significantly prolonged survival in orthotopic GBM xenograft models. These results are consistent with prior studies reporting enhanced therapeutic efficacy of combined MET and mTOR inhibition in other malignancies (46–48), and extend these findings by elucidating a mechanistic basis for synergy in GBM.

Notably, our study reveals a context-dependent shift in the mode of cell death following combined MET and mTOR inhibition. While the combined treatment induces robust apoptosis in untreated parental GICs, crizotinib-pretreated cells preferentially undergo necroptosis upon subsequent mTOR inhibition. This suggests that prolonged MET inhibition primes tumor cells for alternative stress-induced death pathways. Everolimus-mediated disruption of mTOR signaling likely imposes severe metabolic and survival stress. Under these stress conditions, cells may become less capable of initiating apoptosis and subsequently shift to necroptosis as an alternative regulated cell death mechanism. Unlike apoptosis, necroptosis is caspase-independent and is often activated under conditions of overwhelming cellular stress, leading to membrane disruption and inflammatory signaling (49). This phenomenon may represent a unique therapeutic opportunity, as necroptosis could potentially overcome the limitations of apoptosis-driven therapies, particularly in tumor types that exhibit resistance to conventional treatments.

In vivo, we further demonstrate that treatment scheduling critically influences therapeutic outcome. While concurrent administration of crizotinib and everolimus significantly prolongs survival, a sequential regimen—initial MET inhibition followed by mTOR blockade—yields superior survival benefit. This enhanced efficacy may reflect progressive activation and dependency on mTOR signaling during prolonged MET inhibition, effectively creating a state of “mTOR addiction” that sensitizes tumor cells to subsequent mTOR blockade. These findings highlight the importance of therapeutic sequencing in maximizing treatment efficacy and overcoming adaptive resistance in GBM.

Despite these promising findings, several limitations must be considered. The experimental models employed, including GIC cultures and orthotopic xenografts, do not fully recapitulate the complexity of human GBM. In particular, necroptosis is inherently pro-inflammatory and may influence anti-tumor immune responses. Because our models lack an intact immune system, the contribution of immune-mediated mechanisms could not be assessed and should be explored in future studies using immunocompetent systems. Additionally, while our work focuses on MET–mTOR signaling, other compensatory pathways may also contribute to resistance, particularly within the mesenchymal GBM subtype, and merit further investigation.

In summary, our study identifies a previously unrecognized BNIP3–mTOR signaling axis as a key determinant of resistance to MET inhibition in GBM. Dual targeting of MET and mTOR overcomes adaptive resistance, engages multiple cell death pathways, and significantly improves therapeutic efficacy in preclinical models without overt toxicity. These findings provide a strong mechanistic and translational rationale for clinical evaluation of combined MET and mTOR inhibition in patients with GBM, particularly those with high MET expression.

## Author contributions

Y.L., H.K., and I.O. designed the experiments, analyzed the data, and wrote the manuscript with input from all of the authors. Y.L., H.K., S.D., and W.B. performed the experiments. H.V., Y.U., and V.W. performed data analysis. M.M., D.A., M.G., P.G., S.Y., and T.S. provided input. J.L. generated cell lines and provided input.

