## Supplemental Figures for "BNIP3-mTOR Signaling Mediates Resistance to MET Inhibition in Glioblastoma"

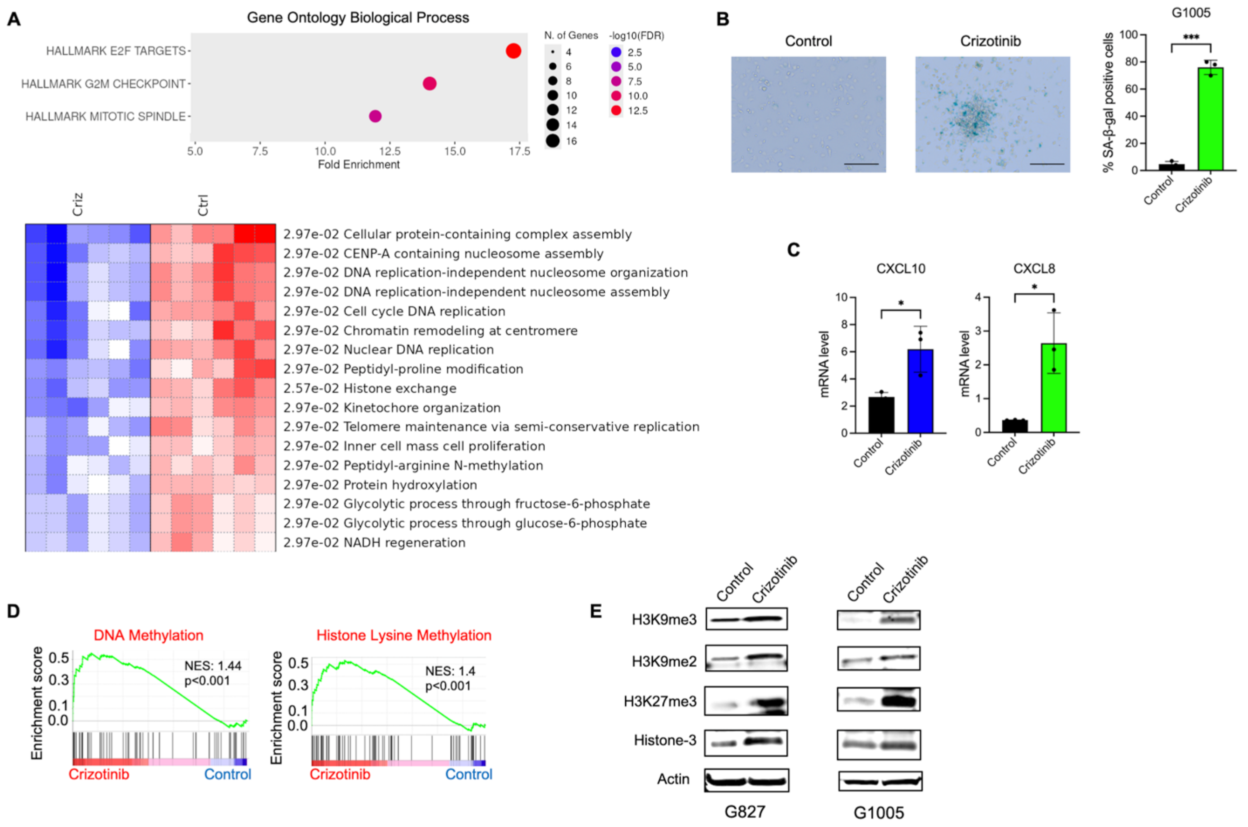


**Supplementary Figure 1: Crizotinib treatment induces transcriptional, epigenetic, and senescence-associated changes in cancer cells.** (A) Gene Ontology (Biological Process) and Hallmark pathway enrichment analysis of RNA-seq data. (B) Representative images of senescence-associated β-galactosidase (SA-β-gal) staining in control and crizotinib-treated GICs, showing increased senescent cells upon MET inhibition. Crizotinib (1.5 µM). Scale bar, 200 µm. Data are shown as mean ± SEM. (****p < 0.0001; two-tailed *t*-test). (C) Quantitative RT–PCR analysis of CXCL10 and CXCL8 mRNA expression in GICs following crizotinib (1.5 µM) treatment. Data represent mean ± SEM. *P* < 0.05 by two-tailed *t*-test. (D) GSEA showing enrichment of DNA methylation– and histone lysine methylation–related gene sets in crizotinib-treated cells compared with control cells. (E) Immunoblot analysis of histone methylation marks in G827 and G1005 cells treated with vehicle or crizotinib.

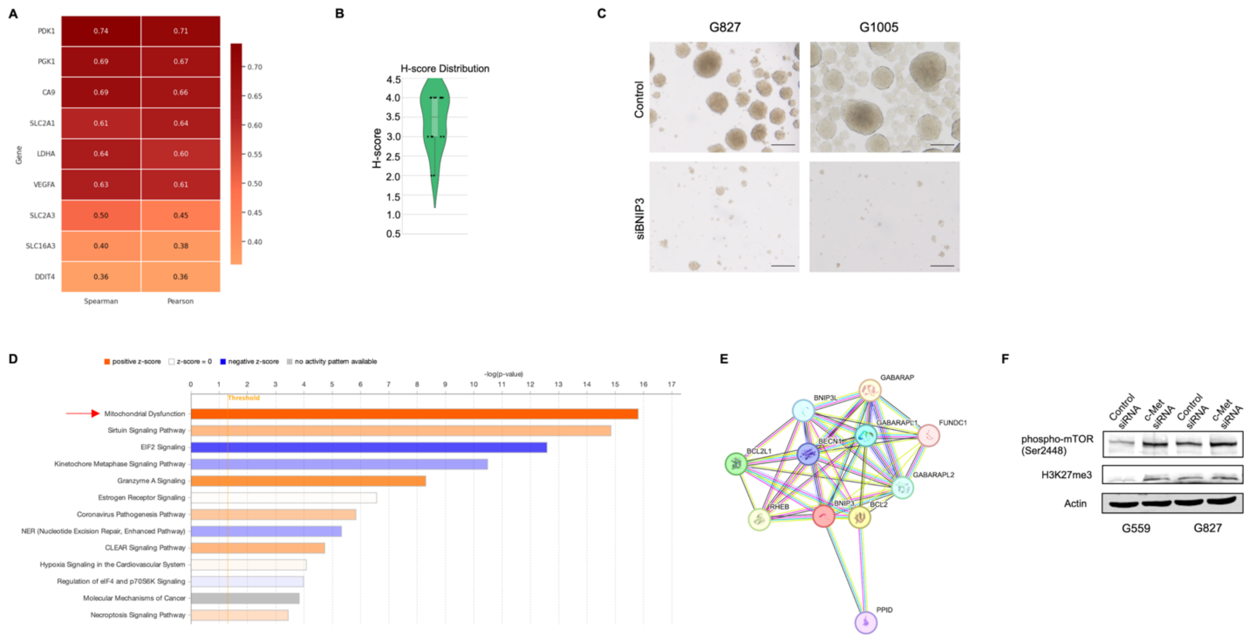

**Supplementary Figure 2: BNIP3 links mitochondrial dysfunction and hypoxia signaling to tumor spheroid growth.** (A) Heatmap showing Spearman and Pearson correlation coefficients between BNIP3 expression and selected HIF1alpha-related genes. (B) Quantification of BNIP3 protein expression using H-score analysis of human GBM samples. n = 28 (from Ivy GBM Atlas Project). (C) 100 GICs were cultured in 24-well plates over one week to compare sphere formation upon transfected with control siRNA or BNIP3-targeting siRNA. Scale bars, 100 µm. (D) Ingenuity Pathway Analysis of differentially expressed genes, highlighting significantly enriched canonical pathways. (E) Protein–protein interaction network of BNIP3. (F) Immunoblot analysis of phospho-mTOR (Ser2448) and H3K27me3 levels in GICs cells transfected with control siRNA or BNIP3-targeting siRNA. Actin was used as a loading control.

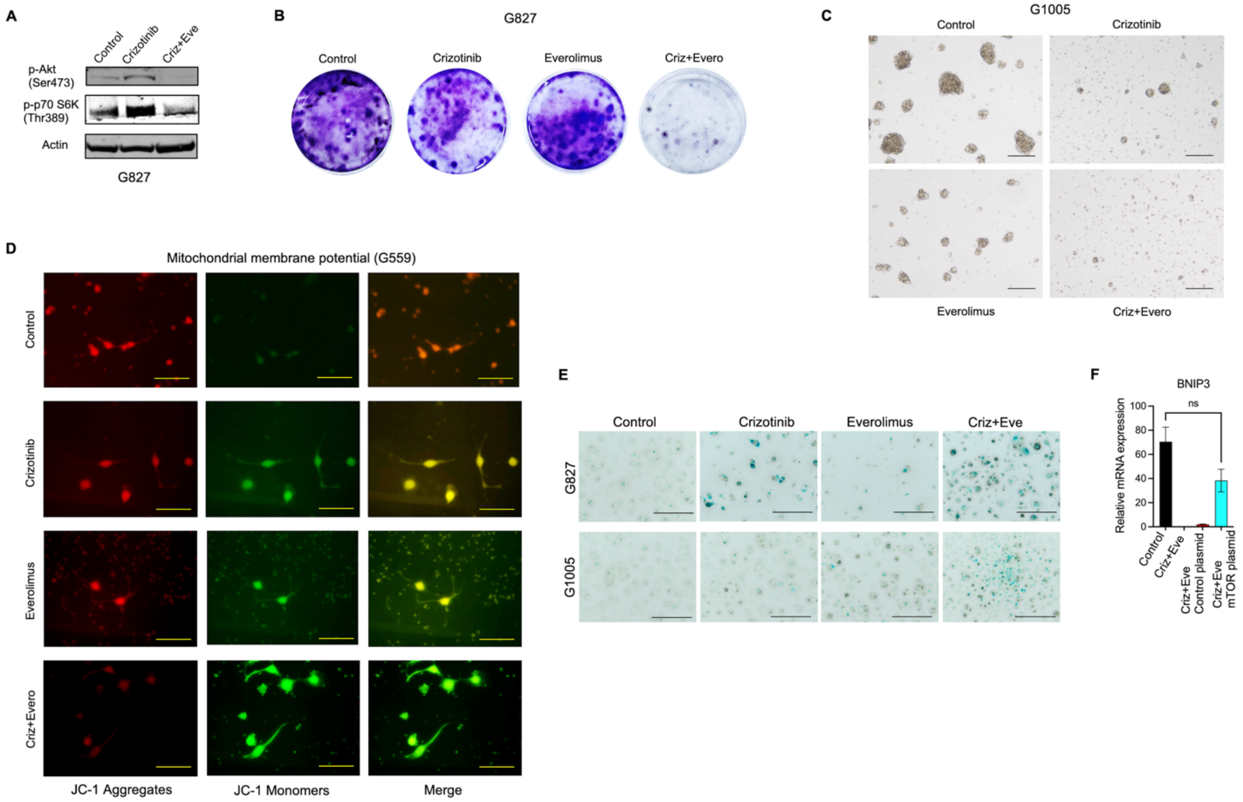


**Supplementary Figure 3: Combined inhibition of MET and mTOR signaling modulates tumor cell growth and mitochondrial function.** (A) Immunoblot analysis in GICs treated with control, crizotinib, everolimus, or their combination (Criz+Evero). (B) Representative colony formation assay images of G827 cells under indicated treatments. (C) 100 GICs were cultured in 24-well plates over one week to compare sphere formation upon treatment with vehicle, crizotinib (1.5 µM), everolimus (5 µM), and the combination of crizotinib and everolimus. Scale bars, 100 µm. (D) Assessment of mitochondrial membrane potential in G559 cells using JC-1 staining. Red fluorescence indicates JC-1 aggregates (intact membrane potential), whereas green fluorescence indicates JC-1 monomers (depolarized mitochondria). Scale bars, 200 µm. (E) Representative images of senescence-associated β-galactosidase staining. Scale bar, 200 µm. (F) Quantitative RT–PCR analysis of BNIP3 mRNA expression under indicated treatments. Data are mean ± SEM ($n$ = 3). One-way ANOVA with Tukey's post hoc test; ns = not significant.
